# Artificial scaffolds that mimic the plant extracellular environment for the culture and attachment of plant cells

**DOI:** 10.1101/2020.06.05.136614

**Authors:** Ryan Calcutt, Richard Vincent, Derrick Dean, Treena Livingston Arinzeh, Ram Dixit

**Author notes:** Corresponding authors Derrick Dean., Treena Arinzeh., Ram Dixit.

## Abstract

Plant growth and development involves an intricate program of cell division and cell expansion to generate different cell types, tissue patterns and organ shapes. Plant cells are stuck together by their cell walls and the spatial context of cells within tissues plays a critical role in cell fate specification and morphogenesis. An *in vitro* model system to study plant development and its regulation by various extrinsic and intrinsic factors requires the ability to mimic the physical interactions between cells and their environment. Here, we present a set of artificial scaffolds to which cultured tobacco BY-2 cells adhere without causing morphological abnormalities. These scaffolds mimic native plant cell walls in terms of their fibrous nature, charge, hydrophobicity and piezoelectricity. We found that the extent of plant cell adhesion was essentially insensitive to the stiffness, fiber dimension, and fiber orientation of the scaffolds, but was affected by the piezoelectric properties of scaffolds where adhesion increased on piezoelectric materials. We also found that the plant cell wall polysaccharide, pectin, is largely responsible for adhesion to scaffolds, analogous to pectin-mediated adhesion of plant cells in tissues. Together, this work establishes biomimetic scaffolds that realistically emulate the plant tissue environment and provide the capability to develop microfluidic devices to study how cell-cell and cell-matrix interactions affect plant developmental pathways.

## INTRODUCTION

Plant growth and development is the outcome of complex interactions between genetic, biochemical and mechanical inputs^1,2^. Similar to animals, cell lineage and positional information both play a critical role in plant cell fate specification and tissue patterning^3^. The physical interaction between plant cells is critical for cell-cell communication and affects the way that cells perceive and respond to their environment^4,5^. Cellular responses can in turn spatiotemporally modify developmental signals to affect neighboring cells, sometimes over considerable distances^6^. Thus, multicellular context and recursive effects between cells and their environment are major determinants of plant development.

Plant cells are encased within a rigid cell wall that provides structural support as well as mechanical and biochemical cues that guide cell morphogenesis^6,7,8,9,10,11^. Since plant cells are cemented together, the cell wall also plays a crucial role in mechanically coupling the growth of adjoining cells within tissues and organs^6,11,12^. The plant cell wall is a fibrous composite material consisting primarily of cable-like cellulose microfibrils embedded within a matrix of hemicellulose and pectin polymers. Cellulose microfibrils consist of crystalline chains of β-1,4 glucan and are the major load-bearing elements of plant cell walls^13,14^. Hemicelluloses are branched polysaccharides that associate with cellulose microfibrils to form a cross-linked network^15^. Pectins are complex, polyanionic polymers that contribute to the porosity and extensibility of the cell wall through crosslinking by borate or Ca^2+^ ions^9,16,17,18^. Pectins are also a major component of the middle lamella that glues the cell walls of adjacent plant cells to provide tissue integrity^19^. The cell wall also contains a small amount of proteins that contribute to wall structure, signaling and wall remodeling^20,21,22^.

In mammalian systems, the development of organ-on-a-chip microfluidic devices that mimic complex three-dimensional cell and tissue configurations has greatly advanced mechanistic analysis of tissue formation and function^23^. These devices rely on appropriate substrates for the attachment, proliferation and differentiation of cells to form the desired tissue. By modifying the physical and chemical properties of the substrates, these devices allow nuanced control of the biochemical and mechanical microenvironment for tissue engineering^24,25^. In plant systems, 3D printed and microfluidic devices have been utilized primarily for long-term monitoring, high-throughput phenotyping and gene expression analyses in intact plants^26,27,28,29^. At the cellular level, microfluidic systems have been used to study the growth and mechanical properties of pollen tubes^30,31^. However, adopting such platforms to study diffusely-growing cells that make up the bulk of the plant body plan has been challenging, primarily because we lack biocompatible substrates to which plant cells adhere to.

In this study, our goal was to fabricate and characterize artificial scaffolds for the adhesion and viable culture of diffusely-growing plant cells as a first step towards engineering functional plant tissues in a microdevice. Here, we report several three-dimensional scaffolds that mimic the fibrous structure, charge, hydrophobicity and piezoelectric properties of natural plant cell walls. We show that cultured tobacco Bright Yellow-2 (BY-2) cells adhere tightly to these scaffolds without causing aberrant cell morphology and that cell adhesion involves pectin.

## RESULTS

### Screening fabricated scaffolds for the attachment of living plant cells

To identify three-dimensional scaffolds for plant cell attachment *in vitro*, we created twenty electrospun matrices that varied in composition, hydrophobicity, fiber diameter, fiber alignment and stiffness (Supplemental Table 1). To determine whether plant cells attach to these scaffolds, we used a suspension culture of tobacco BY-2 cells derived from *Nicotiana tabacum* L. cv. Bright Yellow 2. Tobacco BY-2 cells are widely used as a plant cell model system because of their fast growth rate, ease of live imaging and ability to generate transgenic lines using *Agrobacterium*-mediated stable transformation^32^.

To screen the scaffolds for plant cell attachment, tobacco BY-2 cells four days after subculture were co-incubated with sterilized scaffolds. Tubes were kept on a rotary shaker for four days to allow cell adhesion, followed by a vigorous overnight wash with fresh growth medium to remove unattached cells (Figure 1A). Scaffolds were then stained with the lipophilic styryl dye FM 4-64 to visualize any adhered cells using confocal microscopy. The FM 4-64 dye rapidly labels the plasma membrane of cells and subsequently labels endomembrane compartments due to uptake by endocytosis^33^. Under these screening conditions, we found seven scaffolds that consistently showed adhered BY-2 cells: cellulose acetate (CA), polycaprolactone (PCL), polylactic acid (PLA), and four polyvinylidene-trifluoroethylene (PVDF-TrFE) copolymers (Figure 1B-1H). These scaffolds varied in fiber diameter (nanofiber or microfiber) and orientation (random or aligned) (Figure 3C). Importantly, the attached BY-2 cells were morphologically indistinguishable from control cells grown in the absence of scaffolds. We saw normal-looking BY-2 cell files when stained with the FM 4-64 dye (Figure 1I) and when imaged using an environmental scanning electron microscope (Figure 1J). This is in contrast to a previous study that used scaffolds containing a mix of poly(ethylene terephthalate) (PET) microfibers and PLA nanofibers, which resulted in abnormal plant cell morphology^34^.

**Figure 1.**
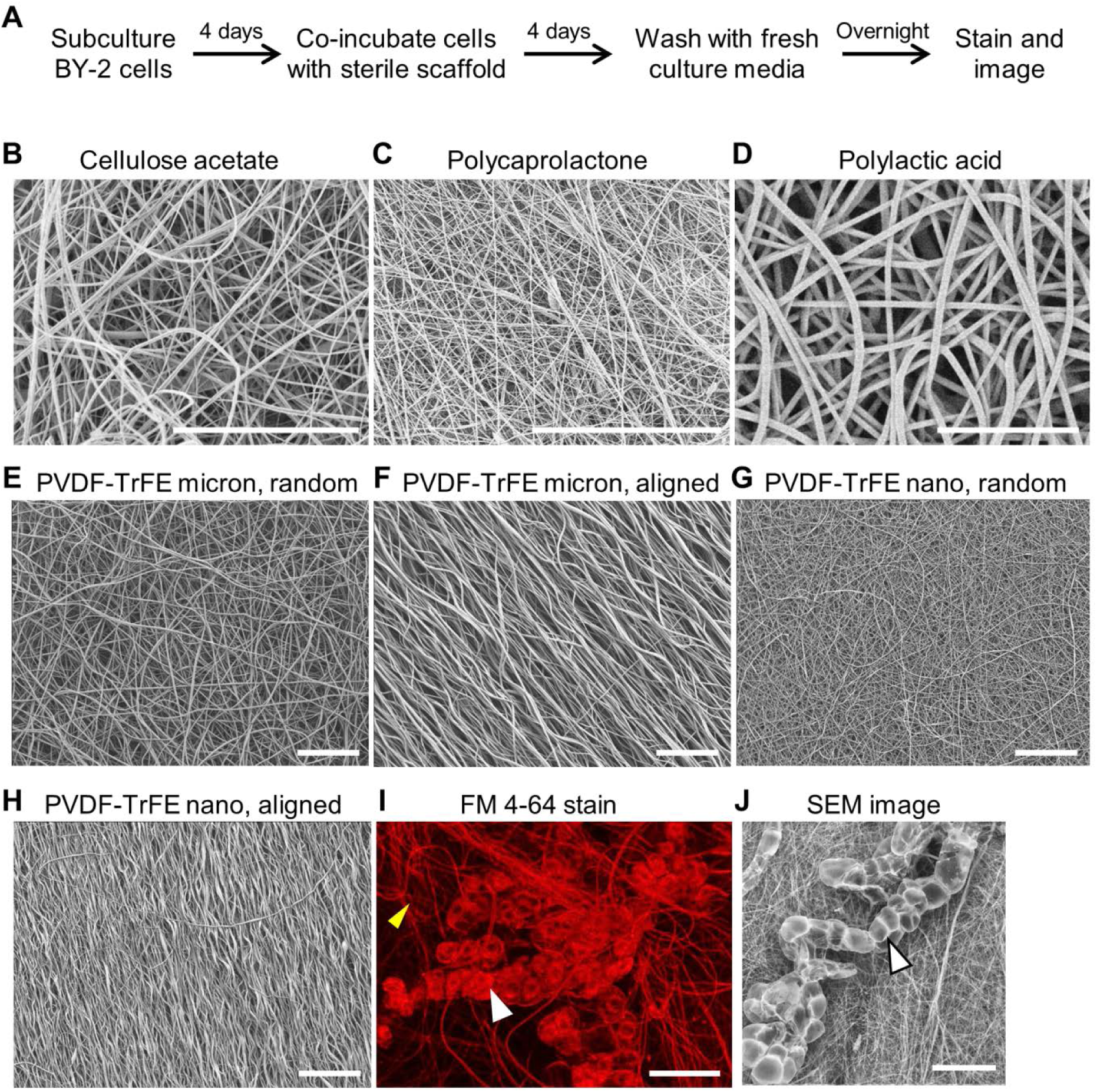
Identifying artificial scaffolds to which plant cells adhere. (A) Outline of the screening procedure used to determine whether cultured tobacco BY-2 cells adhere to electrospun scaffolds. (B-H) Scanning electron micrographs of scaffolds to which BY-2 cells adhere. (I) Fluorescence micrograph of BY-2 cells attached to a microfiber, randomly-oriented PVDF-TrFE scaffold stained with FM 4-64. The white arrowhead points to BY-2 cells while the yellow arrowhead points to scaffold fibers stained by the FM 4-64 dye. (J) Environmental scanning electron micrograph of BY-2 cells attached to a microfiber, randomly-oriented PVDF-TrFE scaffold. Scale bar is 30 µm in (B-D), and 100 µm in (E-J).

To determine which of these seven scaffolds are optimal for plant cell attachment, we sought to quantify the cell density on each type of scaffold. While the FM 4-64 staining revealed attached cells, it was challenging to distinguish individual cells using this dye, particularly in dense fields. In addition, this dye stained the fibers of some of the scaffolds (Figure 1I), further hampering cell quantification. To overcome these problems, we used DAPI (4′,6-diamidino-2-phenylindole) to stain cell nuclei and developed an ImageJ macro to count the number of discrete nuclei on each scaffolds as a proxy for cell abundance (Figure 2A). This quantification revealed that BY-2 cell adhesion varied extensively between the seven scaffolds, with cellulose acetate having the least attached cells and the four PVDF-TrFE scaffolds having the most attached cells (Figure 2B). To determine whether the cells attached to PVDF-TrFE scaffolds are alive, we used the vital dye fluorescein diacetate (FDA). We found that about 82% of adhered cells showed FDA-derived fluorescein signal in the cytoplasm (Figure 2C), indicating that attachment to a PVDF-TrFE scaffold does not induce cell death.

**Figure 2.**
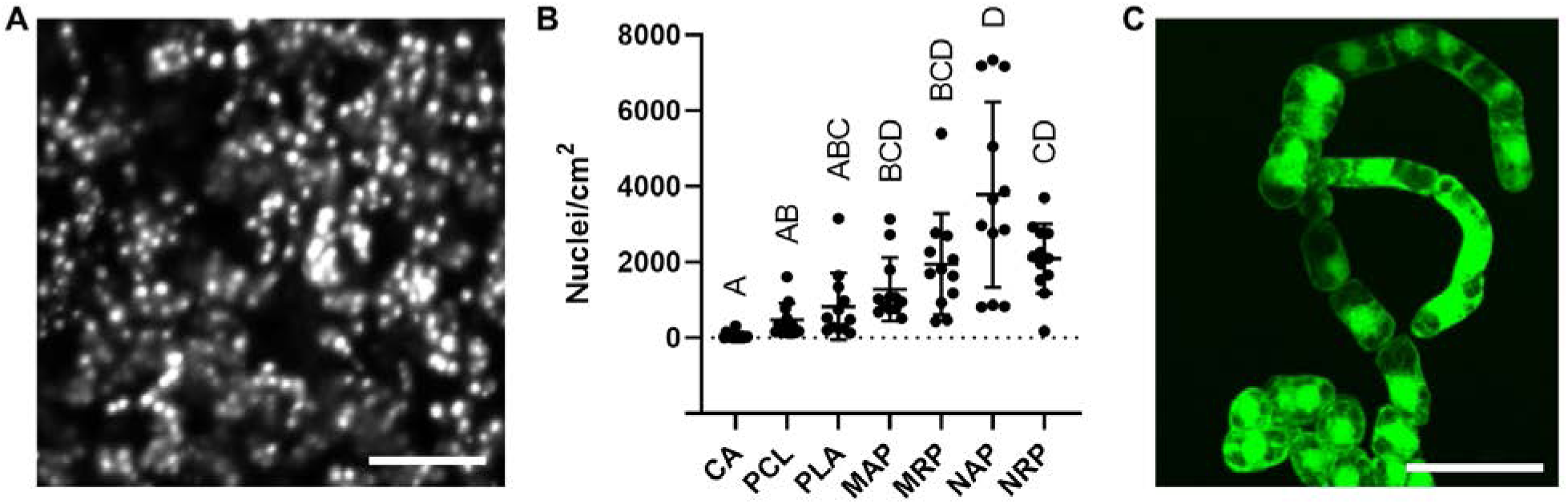
Quantifying the extent of cell adhesion to scaffolds used in this study. (A) Fluorescence micrograph of tobacco BY-2 cell nuclei stained with DAPI on a microfiber, randomly-oriented PVDF-TrFE scaffold. Scale bar = 100 µm. (B) Quantification of cell nuclei found on cellulose acetate (CA), polycaprolactone (PCL), polylactic acid (PLA), microfiber, aligned PVDF-TrFE (MAP), microfiber, randomly-oriented PVDF-TrFE (MRP), nanofiber, aligned PVDF-TrFE (NAP), nanofiber, randomly-oriented PVDF-TrFE (NRP). Center line and error bars indicate mean ± SD from 12 independent biological replicates for each type of matrix. The letters indicate statistically distinguishable groups as determined by the Kruskal-Wallis test with a Dunn’s test for multiple comparisons. (C) Fluorescence micrograph of BY-2 cells stained with fluorescein diacetate on a microfiber, randomly-oriented PVDF-TrFE scaffold. Scale bar = 100 µm.

**Figure 3.**
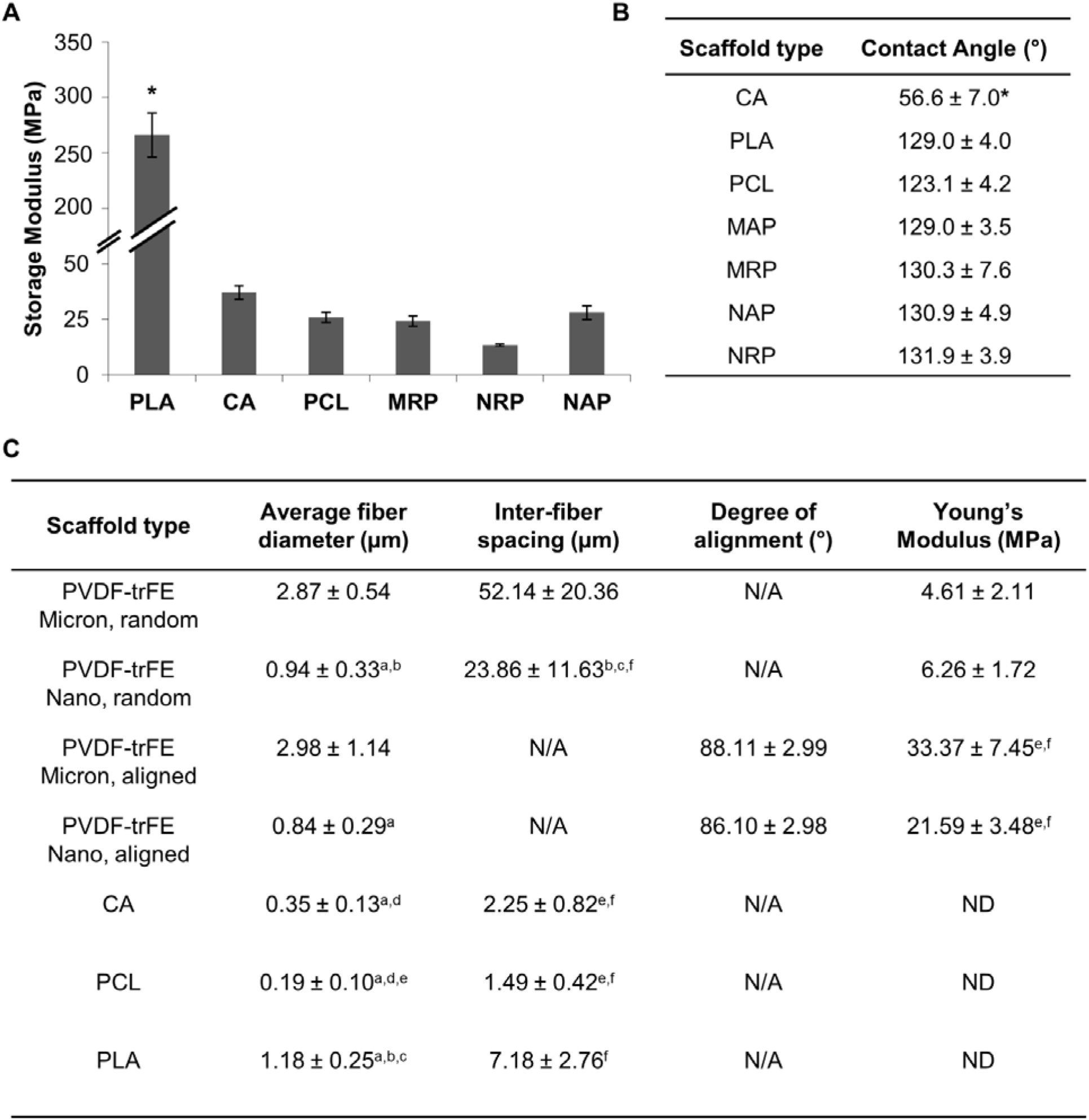
Physical properties of scaffolds used in this study. (A) Storage modulus of scaffolds determined using dynamic mechanical analysis. Values are mean ± SEM. * indicates statistically different from all other groups as determined by one-way ANOVA (p < 0.05). (B) Air-water contact angle at the scaffold surface measured using a goniometer. A contact angle below 90° indicates that the surface is hydrophilic, while a contact angle above 90° indicates that the surface is hydrophobic^47^. Values are mean ± S.D. (n = 4). * indicates statistically different from all other groups as determined by one-way ANOVA (p < 0.05). (C) Table showing the fiber diameter, inter-fiber spacing, fiber alignment and Young’s modulus of the indicated PVDF-TrFE, CA, PCL, and PLA scaffolds. Values are mean ± S.D. ^a^p<0.05, significantly different from PVDF-TrFE Micron group; ^b^p<0.05, significantly different from PCL; ^c^p<0.05, significantly different from CA; ^d^p<0.05, significantly different from PLA; ^e^p<0.05, significantly different from PVDF-TrFE Nano, Random group; ^f^p<0.05, significantly different from PVDF-TrFE Micron, Random group. N/A, not applicable. ND, not determined.

### Characterization of scaffold properties

We wanted to determine whether plant cell attachment correlates with specific properties of the artificial scaffolds. Because mammalian cells are sensitive to scaffold stiffness, we tested whether differences in scaffold stiffness explained the range of BY-2 cell adhesion we observed. Dynamic mechanical analysis was used to measure the storage modulus as an indicator of the stiffness of each scaffold. The storage modulus was measured by applying an oscillating tensile deformation to the samples. The deformation was small (0.1%) to ensure that the testing was conducted in the linear viscoelastic range. The lag in response to this sinusoidal deformation is known as the phase angle and is used to calculate the storage modulus, which is a measure of the elastic response (i.e. energy stored) of the sample. We found that the PLA scaffold had a higher storage modulus than CA, PCL and PVDF-TrFE scaffolds (Figure 3A). Since the CA, PCL and PVDF-TrFE scaffolds have comparable moduli but differ greatly in BY-2 cell attachment, we conclude that scaffold stiffness is not a major determinant of plant cell attachment.

Next, we quantified scaffold hydrophobicity by measuring the air-water contact angle at the matrix surface using a goniometer. This analysis showed that all scaffolds were hydrophobic with the exception of CA (Figure 3B). While hydrophobicity may contribute to cell adhesion, it does not explain the variance between the PCL, PLA, and PVDF-TrFE scaffolds.

At the ultrastructural level, the plant cell wall consists of fibers that vary in dimensions and alignment^35,36^. To examine whether the diameter and spatial organization of scaffold fibers influence the adhesion of plant cells, we took advantage of our PVDF-TrFE scaffolds that have the same chemical composition but differ in fiber diameter and alignment (Figure 3C). We found that while the nanofiber PVDF-TrFE scaffolds had higher mean cell counts than the microfiber counterparts (Figure 2B), these differences were not statistically distinguishable. In addition, there was no significant difference in cell attachment between PVDF-TrFE scaffolds made of randomly-oriented or aligned fibers. Similarly, although the PVDF-TrFE scaffolds with aligned and random fiber arrangements differ in Young’s moduli (Figure 3C), cell attachment was not affected.

### The piezoelectric property of PVDF-TrFE scaffolds contributes to plant cell attachment

A distinctive feature of the PVDF-TrFE scaffolds is that they are piezoelectric^37^, similar to cellulose found in the cell wall^38^. To examine the piezoelectric property’s effect on cell attachment, corona poling was used to enhance the piezoelectric properties of PDVF-TrFE scaffolds (Figure 4A). Corona poling orients the dipoles more uniformly in PVDF-TrFE^39^. Polarization of PVDF-TrFE leads to a positively charged side and a negatively charged side that have significantly lower and higher piezoelectric activity, respectively, compared to an unpoled PVDF-TrFE scaffold as determined by their piezoelectric coefficient, D_33_, values (Figure 4B). Importantly, the average fiber diameter, inter-fiber spacing, Young’s modulus and air-water contact angle of the positive and negative surfaces of poled scaffolds are not significantly different (Figure 4C and 4D). X-ray diffraction analysis showed that while the positively poled PVDF-TrFE has both α-phase (non-piezoelectric) and β-phase (piezoelectric) crystalline structures, the negatively poled PVDF-TrFE has essentially only β-phase (Figure 4E), consistent with its enhanced piezoelectric activity. Interestingly, BY-2 cells cultured with poled PVDF-TrFE scaffolds showed that cells strongly prefer to adhere to the negatively charged side than the positively charged side (Figure 4F).

**Figure 4.**
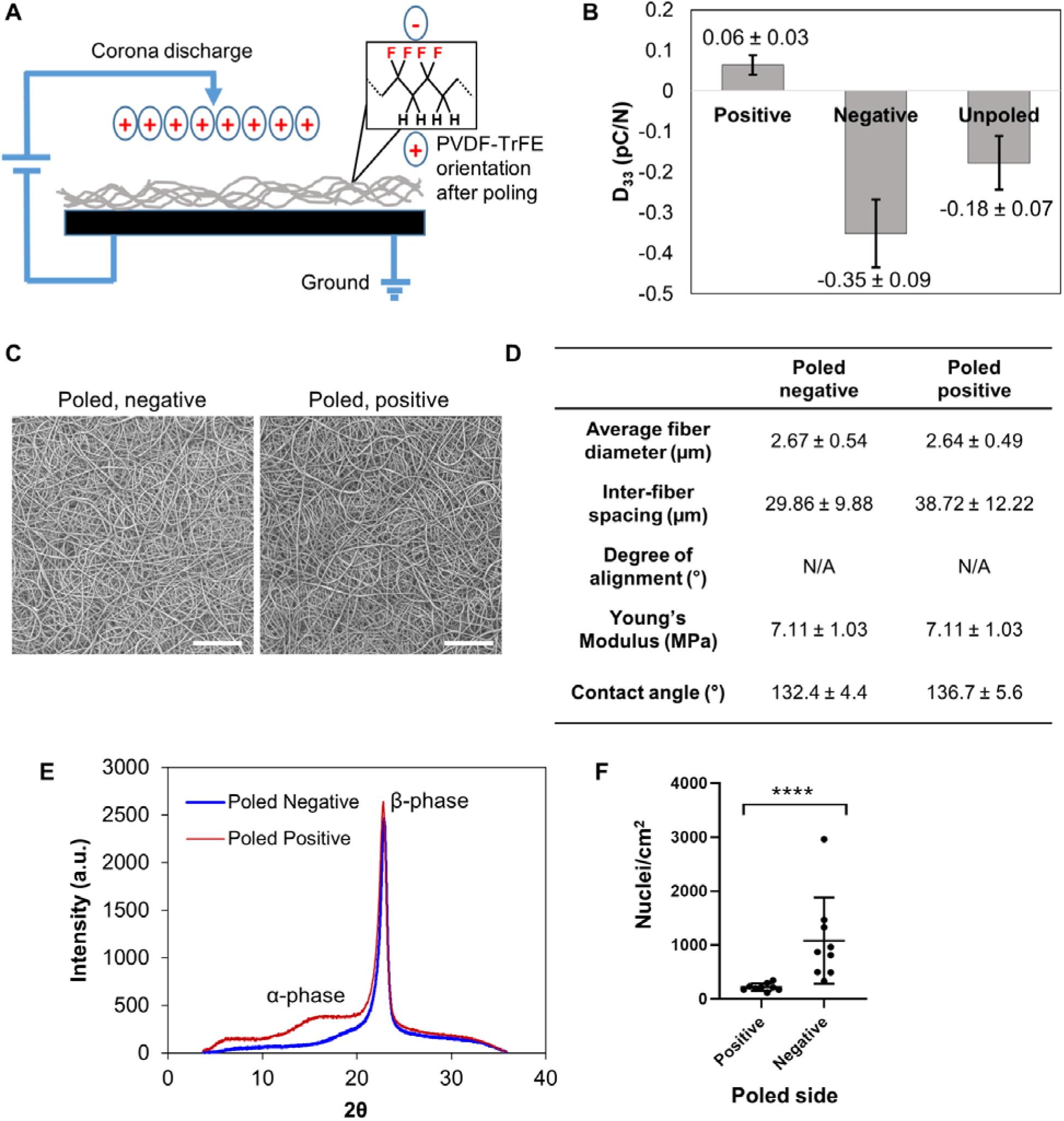
Effect of scaffold piezoelectricity on cell adhesion. (A) Corona poling setup where positive ions are produced in the atmosphere which orients the dipoles in the PVDF-TrFE scaffold resulting in one side being negatively charged and the other side being positively charged. (B) Piezoelectric coefficient (D_33_) of unpoled, poled positive (positively charged surface) and poled negative (negatively charged surface) microfiber, randomly-oriented PVDF-TrFE scaffolds. The sign of the coefficient indicates the surface charge. Values are mean ± S.D. (n = 5). The D_33_ values of the poled positive and poled negative surfaces are significantly different from that of the unpoled surface as determine by one-way ANOVA (p < 0.05). (C) Scanning electron micrographs of poled negative and poled positive PVDF-TrFE surfaces. Scale bar = 100 µm. (D) Table showing the fiber diameter, inter-fiber spacing, fiber alignment, Young’s modulus and air-water contact angle of the poled negative and poled positive PVDF-TrFE surfaces. Values are mean ± S.D. (n = 4 per scaffold). (E) X-ray diffraction characterization of the effect of corona poling on the β-phase structure of the positive and negative surfaces of poled PVDF-TrFE scaffolds. (F) Quantification of cell nuclei found on poled positive and poled negative surfaces. Center line and error bars indicate mean ± S.D. from 9 independent biological replicates for each type of surface. **** indicate p < 0.0001 as determined by the Mann-Whitney test.

### Cell wall pectins mediate the adhesion of plant cells to scaffolds

To determine how BY-2 cells adhere to scaffolds, we treated cells attached to unpoled randomly-oriented nanofiber PVDF-TrFE scaffolds with enzymes that degrade specific components of the cell wall. We measured the abundance of cell nuclei before and after one hour of enzyme treatment to assess whether the treatment detached cells compared to mock-treated scaffolds. Treatment with either 0.05% trypsin or 0.8 % cellulase did not significantly decrease the cell nuclei abundance on scaffolds compared to mock treatment (Figure 5A, 5B and 5D). However, treatment with 0.2% pectinase resulted in an approximately 2-fold decrease in the cell nuclei abundance on scaffolds compared to mock treatment (Figure 5C and 5D). Furthermore, staining of BY-2 cells on PVDF-TrFE scaffolds with the pectin-labeling dye, ruthenium red, showed diffuse red staining surrounding BY-2 cells (Figure 5E), indicating that pectin on the surface of BY-2 cells might facilitate their attachment to scaffolds.

**Figure 5.**
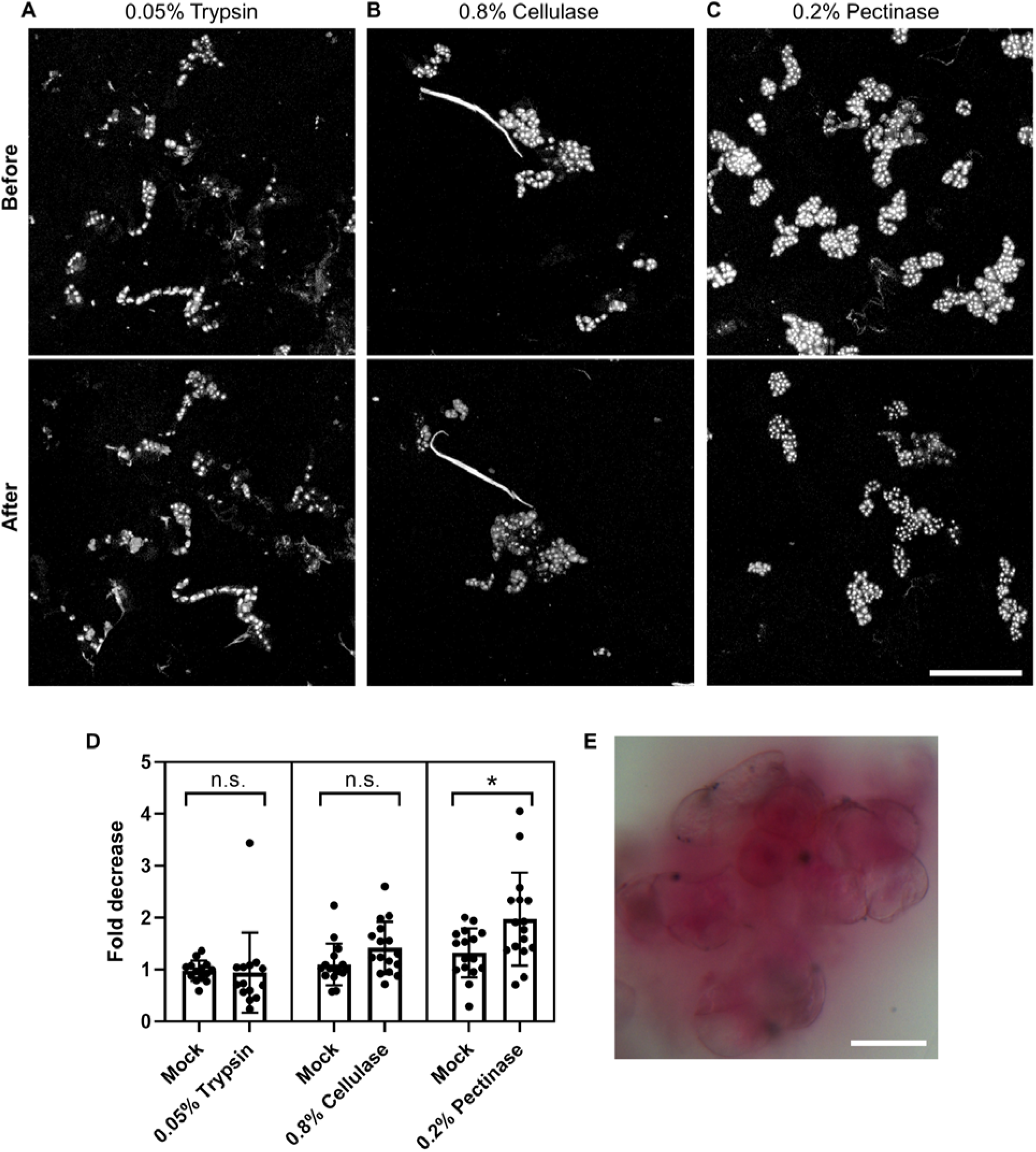
Pectin contributes to attachment of cells to scaffolds. (A,B,C) Fluorescence micrographs of DAPI-stained BY-2 nuclei before and after 1-hr, room temperature treatment with 0.05% trypsin (A), 0.8% cellulase (B), 0.2% pectinase (C). Scale bar = 250 µm. (D) Plot of fold decrease in the abundance of cell nuclei on scaffolds treated with 0.05% trypsin, 0.8% cellulase, and 0.2% pectinase treatments compared to mock treatments (trypsin n = 14, cellulase and pectinase n = 16). * indicates p < 0.05 as determined by the Mann-Whitney test. (E) Pectin staining with ruthenium red of BY-2 cells on a microfiber, randomly-oriented PVDF-TrFE scaffold. Scale bar = 50 µm.

## DISCUSSION

Tissue engineering in mammalian systems was made possible by the development of natural and synthetic matrices that mimic the natural cell environment. These tools greatly accelerated discovery of the fundamental mechanisms for the formation and function of specific tissue constructs and their application for tissue regeneration and repair. Here, we describe the first three-dimensional scaffolds that support the adhesion of plant cells without causing toxicity or altering normal cell morphology. This advance will enable fabrication of plant-on-a-chip devices that will incorporate these scaffolds to study how interactions between plants cells and their environment affect cell behavior, differentiation and tissue patterning.

In our experiments, we tested scaffolds generated using both natural and synthetic materials and with varying material properties. We expected that cells would preferentially adhere to materials derived from plant cell walls, but this was not the case. The best performing scaffolds were synthetic PVDF-TrFE co-polymers and not scaffolds made of cellulose acetate or ones that included cellulose nanocrystals. Therefore, plant cells can adhere to artificial scaffolds, similar to mammalian cells. However, in contrast to mammalian cells, we found that scaffold stiffness, fiber dimensions and fiber orientation are not critical determinants of plant cell adhesion.

All seven scaffolds that supported plant cell adhesion are fibrous materials that mimic the topography of plant cell walls. With the exception of PCL, the other six scaffolds are also negatively charged, similar to the plant cell wall. Intriguingly, six of the seven successful scaffolds are hydrophobic. This is similar to the interaction of plant cells to hydrophobic PET microfiber infused with PLA nanofiber scaffolds^34^. While the physiological relevance of scaffold hydrophobicity is unclear, it is possible that this mimics hydrophobic surfaces of cellulose microfibrils and other cell wall materials^40^.

We found that piezoelectric activity is a key scaffold feature for plant cell attachment. Upon corona poling of PVDF-TrFE, cells almost exclusively attached to the maximally piezoelectric negative surface compared to the positive surface. Piezoelectric activity is a fundamental property of cellulose^41^ and wood^42^, but the implications of this property to plant cells are largely unknown. It is possible that electrical potential created by fiber movement due to fluid flow of the surrounding medium acts as a signal for plant cell adhesion. Alternatively, plant cells might be responsive to an electric charge generated in scaffolds by cell adhesion. In any case, it will be interesting to study whether the piezoelectric activity of cell walls contributes to strengthening of cell-cell adhesion and electrical signaling between cells in growing plant tissues.

Of all the scaffolds we tested, the PVDF-TrFE scaffolds most closely emulated fundamental cell wall properties such as a fibrous structure, negative charge, hydrophobicity and piezoelectricity. Therefore, successful scaffolds for plant cells cannot simply be passive structural materials, but rather they need to mimic the distinctive chemical and physical characteristics of plant cell walls. While animal cells adhere to the extracellular matrix through transmembrane proteins like integrins, plant cells adhere to each other through the pectin-rich middle lamella. Through enzymatic treatments, we determined that pectin, but not proteins or cellulose, plays a major role in adhering plant cells to scaffolds. Consistent with this data, ruthenium red staining demonstrated that BY-2 cells have abundant pectin on their surface. Our findings are in agreement with previous work that indicated that BY-2 cells produce a layer of pectin around four days after subculture^43^. Since the adhesion of BY-2 cells to scaffolds is reminiscent of the way cells adhere in plant tissues, our scaffolds should be suitable for plant tissue engineering. While our data revealed pectin to be important for cell adhesion to scaffolds, we cannot exclude the possibility that other cell wall components such as hemicelluloses also play a role. In addition, additional work is required to uncover the mechanism by which pectins mediate the adhesion of plant cells to scaffolds. Since both the scaffolds and pectins are negatively charged, one possibility is that calcium and/or borate ions might crosslink pectins to scaffold fibers similar to the way these ions mediate pectin crosslinking in the middle lamella^19, 44^. In any case, a pectin-based adhesion mechanism is promising for transferring this technology to other plant cells, since pectin is found in all land plants.

## METHODS

### Fabrication of Scaffolds

Scaffolds were prepared using the electrospinning technique. Random or aligned fibrous PVDF-TrFE scaffolds were fabricated by modifying previous methods as described^45^. Briefly, 17.5% or 20% (w/v) solutions of PVDF-TrFE (70/30) (400 kDa., PDI of 2.1, Solvay Solexis, Inc., Thorofare, NJ) in methyl ethyl ketone (MEK; Sigma-Aldrich, Inc.) was loaded into a syringe with an 18-gauge needle. To generate nano-sized fibers, a voltage of 28 kV was applied to the tip of the needle when electrospinning 17.5% (w/v) PVDF-TrFE solutions. Random fibers were connected on a grounded plate and aligned fibers were collected on a grounded drum at 3200 RPM. The solution was expelled from the syringe at a flow rate of 3 ml/hr at a distance of 40 cm from the collector. Similarly, to generate micron-sized fibers, a voltage of 22.5 kV was applied to the tip of the needle when electrospinning 20% (w/v) PVDF-TrFE solutions. Random fibers were connected on a grounded plate and aligned fibers were collected on a grounded rotating drum at 2600 RPM. The solution was expelled from the syringe at a flow rate of 3.3 ml/hr, 30cm from the collector. After electrospinning the scaffolds were annealed as defined in previous methods^37^.

A horizontal electrospinning setup was used to fabricate the PCL and PLA scaffolds. Solutions (15% w/v) were prepared by weighing the appropriate amount of PCL or PLA pellets and dissolving by stirring in dichloromethane. The resulting solutions were transferred to a 10 mL syringe with a 25½ gauge needle and electrospun at a voltage of 15 kV (using a M826 Gamma High-Voltage Research generator). The feeding rate of the PCL and PLA solutions was set to 1 mL/h using a syringe pump (KD Scientific, Holliston, MA). The polymer solution jet ejected from the tip of the needle was collected at ambient temperature on a grounded static aluminum plate (7.62 cm× 7.62 cm) using a sample-to-film distance of 12 cm. For the CA scaffold, a 25% w/v solution of CA was prepared using a 2:1 mixed solvent of dimethylacetamide and acetone. The solution was electrospun using a voltage of 25 KV and the same target and sample-to-film distance parameters used for PCL and PLA.

### Mechanical Testing

Tensile testing was performed to determine the Young’s modulus and maximum tensile stress of PVDF-TrFE matrices. Tensile tests were performed using a mechanical tester (Instron, MA, USA), as previously described^37^. Briefly, matrices were cut into 70 mm x 10 mm strips. A gauge length of 40 mm was used and the strips were subjected to 10 mm/min displacement rate. All scaffolds were sterilized with 100% ethanol for 20 min and immersed in phosphate-buffered saline for 5 min prior to testing. Sample size of 10 per group was used.

### Rheology

Dynamic mechanical analysis of the scaffolds was conducted using a TA Instruments HR-2 Rheometer. Rectangular samples with a width of 3 mm and length of 12 mm were subjected to an oscillatory tensile deformation ranging from 0.1 to 10 Hz and a strain of 0.1% to ensure that the tests were conducted in the linear range.

### Goniometer measurements

Scaffolds were cut into 1 cm x 1 cm samples (n = 4 per group). Wettability measurements were performed by air-water contact angle using a goniometer built in-house. The scaffolds were mounted onto a glass slide. Water (20 μL) was dropped on the scaffold from a height of 1 cm and the contact angle was recorded using a Sony HDR-XR260V camera.

### X-Ray Diffraction

X-ray diffraction scans were collected using a Bruker D8 Discover diffractometer equipped with a general area detection system and Co radiation generated at 40 KV and 35 mA and a collection time of 10 minutes.

### Corona Poling of the PVDF-TrFE Scaffolds

Corona poling was performed using a custom setup. Depending upon the thickness of the matrix used, the PVDF-TrFE matrices was heated using a heating plate to a temperature of 47-55°C for 30-60 min. A metal grid connected to a primary voltage source (set to 2 kV) was suspended 2 cm above the heated matrix. A needle connected to a secondary voltage (set to 14kV) was placed above the grid with the needle tip oriented in the direction of the grid. Charge was discharged for 3-6 hours.

### Measuring Piezoelectric Coefficient (D_33_) of PVDF-TrFE Scaffolds

Scaffolds were cut into 1 cm x 1 cm samples from each group (n = 5 per group). D_33_ values were measured using a piezometer (Piezotest, D_33_ Piezometer System, Singapore). Each matrix was placed onto a piezometer and the D_33_ value measured at a dynamic force of 0.25 N at 110 Hz.

### SEM of Scaffolds

Scaffolds were sputter coated with gold-palladium (EMS 150 TES, PA, USA) and viewed using Scanning Electron Microscope (SEM, JSM-7900F JEOL, USA) with an accelerating voltage of 2-3 kV. Two images in three different areas of each scaffold were captured. Fiber diameters and inter-fiber spacing were measured at n = 15 per image using ImageJ software, as previously described^46^. Degree of alignment for aligned fiber matrices was measured as previously described^46^. Briefly, a line was drawn perpendicular to a fiber, chosen at random, in the SEM image. Ten angles were measured between the line and the fibers. The absolute deviation value (ADV) of the measured angles from 90° was calculated and averaged. The percent alignment was calculated as:

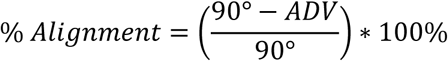

### BY-2 Cell Culture

Tobacco BY-2 cell cultures were maintained in 100 mL of liquid medium (4.3 g/L Murashige and Skoog basal salts, 100 mg/L myo-inositol, 1 mg/L thiamine, 0.2 mg/L 2,4-D, 255 mg/L KH_2_PO_4_ and 3% sucrose, pH 5.7) in 500 mL VWR polycarbonate Erlenmeyer flasks at 25°C on a rotary platform shaker at 130 rpm. Cells were subcultured weekly using a ratio of 1:20 of cell culture to fresh liquid medium.

### Screening Scaffolds for Cell Attachment

Scaffolds were cut into small pieces (approximately 0.75cm × 1cm) and sterilized by submerging in 100% ethanol for 1 hour with shaking to remove air bubbles from the scaffolds. Scaffolds were then dried in a laminar flow hood and rinsed thrice with sterile ultrapure water and once with fresh BY-2 culture medium. Sterile scaffolds were co-incubated with four-day-old BY-2 cell cultures in screw-cap cryotubes and placed on a nutator for 4 days at 50 rpm, then washed with 25 mL of fresh media in a 50 mL centrifuge tube overnight on a nutator at 50 rpm.

### Imaging and Image Analysis

Fluorescence microscopy was performed on a Nikon A1si confocal microscope. For cell counting, cells were stained with 1 µg/mL DAPI (AnaSpec Inc 83210) in BY-2 media for 10 minutes, briefly rinsed in fresh media, and then imaged with a 403 nm diode-pumped solid-state laser (Melles Griot) and 425-475 nm emission filter. For initial screening and cell viability staining, cells were stained with 3 µg/mL FM4-64 (Tocris 5118) in BY-2 media then imaged with a 560 nm diode-pumped solid-state laser (Melles Griot) and 575-620 emission filter. For live/dead cell staining, cells were stained in 2.5 µg/mL fluorescein diacetate (Sigma F7378) for 15 minutes and imaged using a 488 nm Argon-ion laser (Melles Griot) and a 500-550 nm emission filter.

All images were processed using ImageJ. Nuclei counts were obtained using an ImageJ macro that took maximum intensity projections, converted them to 8-bit, and then used the contrast method of the Auto Local Threshold function (radius 50) to create thresholded images of nuclei. The macro then ran Analyze Particles with a 20-200 size filter to count the number of nuclei. To determine the effect of enzyme treatments on cell adhesion, all images were converted to maximum intensity projections, then combined into a single stack for uniform processing. The intensities of the images in the stack were then normalized using the Enhance Contrast function (0% saturation). The integrated signal intensities of the images were measured and used to calculate the fold decrease by dividing the before treatment intensity by the after treatment intensity.

### Enzyme Treatments

Cell nuclei on nanofiber, randomly-oriented PVDF-TrFE scaffolds were imaged after staining for 10 minutes with 1 µg/mL DAPI, then placed in a 1.7 mL microcentrifuge tube with 1 mL of BY-2 media containing either 0.05% trypsin (Sigma T2601), 0.8% cellulase (Duchefa Biochemie C8001), or 0.2% macerozyme (pectinase, Duchefa Biochemie M8002), and shaken at room temperature for one hour. Mock treatments used BY-2 media alone. Scaffolds were then washed in 25 mL of fresh media three times and then staining for 10 minutes with DAPI for imaging.

### Environmental Scanning Electron Microscopy

Scaffolds were co-incubated with BY-2 cells and washed according to our standard protocol, then dried on one side with a Kimwipe and adhered to a mount and imaged using a Zeiss Evo10 environmental scanning electron microscope. Vacuum-chamber vapor point varied between 80-100 Pa and electron high tension value varied between 16 and 20 kV.

### Statistical Analysis

Statistical analyses of nuclear counts and signal intensity changes were performed using GraphPad Prism 8 (GraphPad Software, San Diego, CA, USA). Normality was determined using Anderson-Darling, D’Agostino and Pearson, Shapiro-Wilk, and Kolmogorov-Smirnov tests. For non-normal datasets, the Kruskal-Wallis test with a Dunn’s test was used to determine statistical significance (p < 0.05) for multiple comparisons and the Mann-Whitney test was used for comparison between two groups. For quantitative materials characterization data, statistical analyses were performed in SPSS Statistics Version 25 (IBM, Armonk, NY, USA). One-way ANOVA was used to determine statistical significance (p < 0.05). Normality was determined using the Shapiro–Wilk Test and Levene’s equal variance test. Tukey’s posthoc test was used for statistical differences at p < 0.05. Values are reported as mean ± standard deviation, unless otherwise mentioned.

## ACKNOWLEDGEMENTS

We thank Hanxiao Huang for help with scaffold characterization. This work was supported by the Center for Engineering Mechanobiology (CEMB), an NSF Science and Technology Center, under grant agreement CMMI: 15-48571.

