## Supplemental Table 1 for "Artificial scaffolds that mimic the plant extracellular environment for the culture and attachment of plant cells"

**Supplemental Table 1: Scaffolds tested for plant cell adhesion**

| Scaffold type | Cell adhesion |
|---|---|
| Polyvinylidene-trifluoroethylene (PVDF-TrFE) microfiber, aligned | Yes |
| PVDF-TrFE nanofiber, aligned | Yes |
| PVDF-TrFE microfiber, random | Yes |
| PVDF-TrFE nanofiber, random | Yes |
| 0.8% polylactic acid (PLA) | Yes |
| Polycaprolactone (PCL), loose fibers | Yes |
| Cellulose acetate (CA), random | Yes |
| Polyvinyl alcohol (PVA) | No |
| PCL, compact fibers | No |
| 3D-printed PLA disk coated with electrospun PVA nanofibers | No |
| 3D-printed PLA disk coated with electrospun PVA nanofibers containing 1% cellulose nanocrystals | No |
| PLA + 2.5% cellulose nanocrystals | No |
| PCL + 2.5% cellulose nanocrystals | No |
| PVA + 2.5% cellulose nanocrystals | No |
| Cellulose acetate collector | No |
| Cellulose acetate grid | No |
| Gelatin microfiber, aligned | No |
| Gelatin nanofiber, aligned | No |
| Gelatin microfiber, random | No |
| Gelatin nanofiber, random | No |
